# Diversity and structure of the deep-sea sponge microbiome in the equatorial Atlantic Ocean

**DOI:** 10.1101/2024.01.24.577104

**Authors:** Sam E. Williams, Gilda Varliero, Miguel Lurgi, Jem Stach, Paul R. Race, Paul Curnow

## Abstract

Sponges (phylum Porifera) harbour specific microbial communities that drive the ecology and evolution of the host. Understanding the structure and dynamics of these communities is emerging as a primary focus in marine microbial ecology research. Much of the work to date has focused on sponges from warm and shallow coastal waters, while sponges from the deep ocean remain less well-studied. Here, we present a metataxonomic analysis of the microbial consortia associated with 23 deep-sea sponges. We identify a high abundance of archaea relative to bacteria across these communities, with certain sponge microbiomes comprising more than 90% archaea. Specifically, the archaeal family *Nitrosopumilaceae* are prolific, comprising over 99% of all archaeal reads. Our analysis revealed sponge microbial communities mirror the host sponge phylogeny, indicating a key role for host taxonomy in defining microbiome composition. Our work confirms the contribution of both evolutionary and environmental processes to the composition of microbial communities in deep-sea sponges.

**Importance:** The deep ocean is the largest biome on Earth, accounting for >90% of the planet’s marine environment. Despite this it remains a largely unexplored ecosystem, with less than 0.01% of the deep seafloor having been quantitatively sampled. Deep-sea sponges are ancient metazoans which harbour complex microbial communities and much still remains to be learned about the composition and diversity of these unique microbiomes. In an effort to address this, here we report a metataxonomic analysis of the microbial consortia associated with 23 deep-sea sponges from the equatorial Atlantic Ocean. Our findings reveal intricate, species-specific microbial communities dominated by ammonia-oxidizing archaea. This study highlights the significant role sponges play in shaping microbial consortia, providing new insights into deep-sea ecosystem dynamics. Importantly, our findings provide a scientific basis for understanding the evolutionary relationships between sponges and their symbiotic microorganisms.

## Introduction

Sea sponges (phylum Porifera) live in intimate symbiotic relationships with complex microbial communities (1-3) Microbial abundance within sponge tissue can be orders of magnitude greater than the surrounding seawater (4). Sponge microbiotas are generally dominated by the Pseudomonadota and a few other core phyla (5), but at least 30 other variable phyla are commonly identified in studies of metataxonomic communities. These communities exhibit unusual features such as the presence of the sponge-associated phyla known as Poribacteria, which are only rarely observed living outside the sponge hosts (6, 7). Large-scale comparative surveys have confirmed that microbial communities are relatively stable within, but vary greatly in richness and diversity across, sponge species (6-9). This species dependence suggests that particular microorganisms might be selected for by sponge-microbe or microbe-microbe interactions. However, these relationships might be modulated by other biotic and abiotic factors; for example, water temperature appears to have a major influence on microbial community structure inside hosts (7, 10).

Much of our current knowledge of sponge-associated microbes is derived from samples collected in relatively shallow coastal waters (5). There are fewer studies investigating sponges which inhabit the deep ocean (11), at least in part due to the difficulty in obtaining such samples. Sponge communities from mesopelagic and abyssal depths might be expected to differ from their shallow-water relatives given the drastically different ambient conditions - cold, dark, nutrient-poor - and the consequent changes to the microbial life in the seawater around the sponge. Current evidence suggests that deep-water sponge-associated bacterial communities remain species-specific, and that sponges with both High-and Low Microbial Abundance (HMA and LMA, respectively) are found at these depths (8, 12, 13). Perhaps the major differences in the community structure of deep-sea sponges are the general loss of Cyanobacteriota and a lower occurrence of Poribacteria, as well as the increased abundance of Archaea (12-14). Steinert and colleagues also showed that Hexactinellid glass sponges, which commonly inhabit deep and cold waters, had lower overall microbiome diversity than deep-water demosponges and that there were differences between the bacteria associated with the two classes (12). Archaea are found to be more abundant than bacteria in some deep-water sponges and are almost entirely dominated by the phylum Thermoproteota, specifically family *Nitrosopumilaceae* (14, 15). Only two other archaeal phyla appear to be present (Methanobacteriota, Nanoarchaeota), all at low abundance and without an obvious relationship to sponge species or class (12-14). As with shallow-water sponges, the precise composition of deep-water sponge communities reflects the complex relationship between evolutionary and environmental factors (8, 16).

The recent shift towards whole genome-based taxonomy has significantly revised the microbial tree of life (17). Traditional taxonomic assignment via outdated 16S rRNA databases fail to account for this revision and perform poorly in standardised tests (18). The newly developed, Greengenes2, unifies whole genome taxonomy and 16S rRNA, offers improved accurate taxonomic assignment and has not yet been applied to the sponge microbiome (19). As well as their fundamental role in sponge ecology, the microbial communities of deep-sea sponges are also of increasing interest as a resource for novel therapeutics. Sponge symbionts are well-established as a fertile repository of bioactive natural products (20) and the diverse composition of sponge microbiota has the potential for high and novel functional diversity (21). The microbial consortia of deep-sea sponges remain largely untapped in such biodiscovery programmes, where metataxonomic data collected for community analysis could guide future drug discovery programmes (22, 23).

In this study, we investigate the microbiome of 23 diverse sponges from 5 underexplored deep-water seamounts in the Atlantic Ocean. We focus on elucidating the impact of host-sponge class on microbial composition and use updated taxonomic assignments to better profile these communities. We investigate factors shaping these microbial communities, including the biogeographical factors, and sponge host phylogeny on microbiome composition. Specifically, we explore the extent to which sponge host phylogeny correlates with microbiome structure. This study adds novel insight into host phylogeny’s impact on the sponge microbiome and explores previously unstudied sponge taxa.

## Materials and Methods

### Sample Selection

A subset of 23 deep-sea sponges were chosen from a larger collection which was described previously (24). Sponges were chosen to represent a variety of depths and sampling sites. All the chosen samples were collected between depths of 569-2618 m (Average depth: 1455 m) from five different sampling sites in the Atlantic Ocean by the NERC research vessel RRS James Cook during research cruise JC094 (25) (Figure 1). Samples were provided to us directly by the cruise Chief Scientist Laura Robinson, University of Bristol. Sample handling and preparation was performed according to the protocols adopted for the Earth Microbiome Project (EMP) (26) and subsequently the Sponge Microbiome Project (5). Local sea water controls were not available for sequencing and so are absent from the dataset.

**Figure 1.**
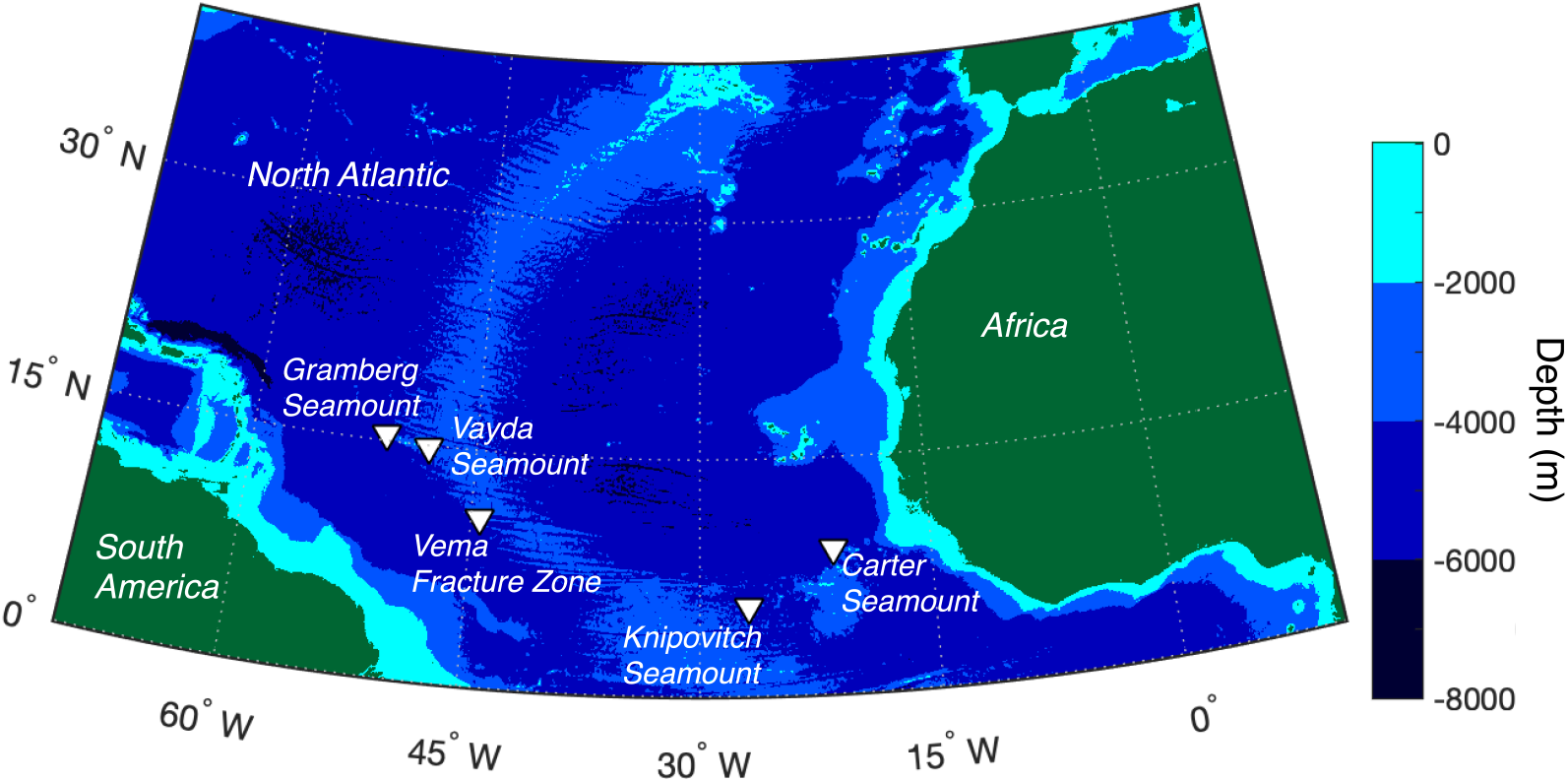
Map of the five deep-sea sampling areas (triangles) from the JC094 research cruise in the mid-Atlantic Ocean. Graphic created using ETOPO1 bathymetry.

### DNA extraction

DNA was extracted from 0.25 g of sponge tissue using the DNeasy PowerSoil Kit (Qiagen, Hilden, Germany) using the optimized procedure of Marotz et al. (27) and including a blank extraction to account for intrinsic kit or reagent contamination (28). Prior to extraction, sponge samples were washed three times in sterile artificial sea water (ASW; Crystal Sea Marine Mix, Marine Enterprise International). All extractions were performed in a laminar flow hood.

### Identification of sponge species

Sponge taxonomy was assigned based upon the mitochondrial cytochrome oxidase subunit I (COI) or the 28S rRNA gene. The COI gene was amplified through PCR using the universal primers LCO1490 and HCO2198 (Supplementary Table 1) (29). The reaction comprised 20 µL Platinum™ Hot Start PCR Master Mix (2X) (Invitrogen, Thermo Fisher Scientific, Waltham, MA, USA), 1.6 µL of each primer at 10 pmol/µl, 14.8 µL deionised water and 2 µL DNA template. Thermal cycling was performed as described by Yang et al. (30): 1 minute denaturation at 94 °C; 5 cycles of 94 °C for 30 sec, annealing at 45 °C for 90 sec and extension at 72 °C for 1 min; 35 cycles of 94 °C for 30 sec, 51 °C for 40 sec and 72 °C for 1 min; and a final extension step at 72 °C for 5 min. 5 µL of the amplified PCR fragment was analysed on a 1% TAE agarose gel to confirm the expected COI gene fragment size of approximately 680bp. Samples unable to be identified by the COI locus were reanalysed with 28S primers NL4F and NL4R (Supplementary Table 1) (31), using the same PCR reagents but with thermal cycling as follows: 10-min initial denaturation at 95 °C; 35 cycles of: 95 °C, 56 °C and 72 °C for 1 minute each; and a final extension step of 72 °C for 7 minutes (30). 5 µL of these reactions were analysed on a 1% TAE agarose gel to confirm 28S gene fragment size 858 to 1010 bp. PCR products were then either purified directly using Monarch PCR and DNA Cleanup Kit (New England Biolabs, Hitchin, UK), or if more than one band was present extracted from 1% TAE gel using the QIAquick Gel Extraction Kit (Qiagen, Hilden, Germany). Nucleotide BLAST (BLASTn) was used for taxonomic assignment (32). Sponge taxonomy was assigned at the species level if BLASTn % identity >97%, we recognise this threshold is arbitrary but sufficient for the purposes of this study (33). For identities below this, assignment was made to the nearest genus, recognizing that lower % identities yield approximate, not precise, taxonomic classifications. To distinguish between sponges of the same species, or unassigned sponges the original sample identifier (BXXXX) was added as a suffix.

### Sponge Phylogenetic Analysis

COI gene sequences were aligned with MAFFT (v7.490) using the L-INS-I algorithm (34). A phylogenetic tree was then constructed using the IQ-TREE2 (v2.2.2.7) with model finder, the best-fitting model was TIM2+F+I+G4 and the analysis included 1,000 ultra-fast bootstrap replicates (35). The trimmed COI gene from the slime sponge *Oscarella carmela* (GenBank: NC_009090.1) was used as an outgroup on which the tree was rooted. The tree was visualised and annotated with iTOL v6 (36).

### Sponge microbiome sequencing

The V4 and V5 regions of the microbial 16S rRNA gene were amplified via PCR with the universal primers 515F-926R (26) combined with Illumina sequencing adaptors (Supplementary Table 1). Thermal cycling followed the standardised protocol of the Earth Microbiome Project (26). This entailed an initial denaturation at 94 °C for 3 minutes, followed by 35 cycles of 94 °C for 45 seconds, 50 °C for 60 seconds and 72 °C for 90 seconds, with a final extension of 10 minutes at 72 °C. The presence of the expected PCR products was confirmed by visualising 5 µL of the reaction on a 1% agarose gel. For most sponge samples this produced a single intense band at approximately 478 bp, and the amplicon pool was cleaned with the QIAquick PCR Purification Kit (Qiagen 28104). If spurious bands were identified the 478 bp fragment was extracted using the QIAquick Gel Extraction Kit (Qiagen 28704) alongside a blank gel lane extraction as a further negative control for environmental contamination.

Illumina sequencing was performed by the genomics facility at the University of Bristol. After initial quality assessment and normalisation of amplicon libraries, samples were indexed with the XT Index Kit (Nextera®) and sequenced using the MiSeq platform (Illumina). Base calling and quality assessment was performed with Real Time Analysis version 1.18.54.0. At least 200,000 reads were recovered for each sample.

### Sponge microbiome sequencing and bioinformatic analyses

Microbiome analysis was performed with QIIME2 2023.7 (37) and the data processing pipeline is available at https://github.com/Sam-Will/Deep_Sponge_Micro. Adaptors and primers were removed from the demultiplexed sequence data using the cutadapt plugin (38) and the subsequent data was denoised with DADA2 (v. 1.26) (39) via the denoised-pair command. Truncation parameters were chosen based on the quality plots to ensure a minimum of 30 bp overlap between reads. Amplicon sequence variants (ASVs) were assigned taxonomy using classify-sklearn feature-classifier (40) trained against the Greengenes2 2022.10 full length sequences classifier (19). Data was imported to R version 4.0.2 for further analysis, and the following packages were used (41). Inherent laboratory contamination during sample handling and eDNA extraction was assessed by analysing blank samples from the DNA extraction kit (350,173 sequences) and from the gel extraction kit (398 sequences). Twenty-six of the ASVs in these negative controls were identified as likely true contaminants by the decontam 1.8.0 package in combined mode (42). These sequences were removed from the sponge dataset (Supplementary Table 2). Identified contaminants were removed from all samples in the data set prior to further analysis. Phylogenies were constructed with phyloseq 1.42.0 and ASV alpha diversity values were calculated using the ‘estimate_richness()’ function and assessed with aov() (41). Rarefaction curves were calculated with iNEXT (v 2.0.20) (43). Permutational multivariate analyses of variance (PERMANOVA), non-metric multidimensional scaling (NMDS) and the distance decay analyses were calculated using Vegan (v 2.5.7) (44). Before performing PERMANOVA, the dataset was tested for homoscedasticity using the function betadisper() and then anova(). As homoscedasticity was observed (p > 0.05 in ANOVA test), PERMANOVA was then performed on the Bray-Curtis dissimilarity matrix calculated on the Hellinger transformed ASV dataset (1000 permutations). PERMANOVA was performed using all samples except from sample Hexactinellid B01175 which was the only representative of >2500 m depth. To calculate the community correlation with sampling depth (i.e., distance decay) and geographical distances, mantel statistical tests (“method = Spearman”) were performed on the Bray-Curtis matrix computed on Hellinger transformed ASV dataset and Euclidean matrix calculated on depth data, and geographical distances (1000 permutations). *p*-values obtained from the Mantel tests were corrected using the Holm-Bonferroni method (45). Geographical distances were calculated using the R library geosphere (v 1.5-1.8) (46). NMDS was performed on the Hellinger transformed ASV dataset. Plots and data manipulation were performed using gplots (v 3.1.1), RColorBrewer (v 1.1.2), tidyr (v 1.1.2), ggplot2 (v 3.3.3) and svglite (v 2.1.2). A dendrogram of the sponge microbiome community structure was obtained using the hclust function (“method = ward.D2”) and was formatted using ape (v 5.7-1) and TreeTools (v 1.10.0) packages (47, 48). All of the above steps were done in the R environment (49). Dendroscope (v 3.8.10) (50) was to create a tanglegram depicting similarities between the microbiome community structure and sponge phylogeny.

### Data Availability

The COI sequences of sponges in this study have been deposited in GenBank assigned the accession numbers PP092504-PP092519 and OP036683.1. All raw 16S rRNA gene amplicon sequencing data is available in the NCBI Sequence Read Archive, associated with BioProject number PRJNA702029. Additionally, the data processing pipeline, including Qiime2 taxonomy, representative sequences, phylogenetic trees, and metadata, can be found at https://github.com/Sam-Will/Deep_Sponge_Micro.

## Results

### Classification of sponge samples

Sponge taxonomy was initially based upon the gene for mitochondrial cytochrome oxidase (29). This approach was used to successfully classify 17 of the 23 sponges considered here at the genus or species level (Figure 2). Assignments for certain samples to the nearest genus, including *Euplectella, Desmacella*, and *Aspidoscopulia,* are tentative since they have a low BLAST identity to their nearest relative and thus are likely novel. Of the remaining 6 sponges, 2 were subsequently identified to species level by the 28S rRNA gene (*Hertwigia* sp. MD-2008 97.09% and *Trachycladus* sp. NCI325 97.51%). The four remaining sponges were identified only to the class level as Hexactinellid, based on a visual inspection of spicules. Overall, the 23 deep-sea sponges studied here comprise 13 demosponges with representatives from 6 orders, 8 families and 10 species; and 10 hexactinellids from 2 orders, 3 families and 5 species (Figure 2, Supplementary Table 3). To our knowledge this dataset comprises several sponge genera whose microbiota has not previously been characterised, including *Rhabderemia, Trachycladus, Euplectella* and *Aspidoscopulia* (11, 12, 51).

**Figure 2.**
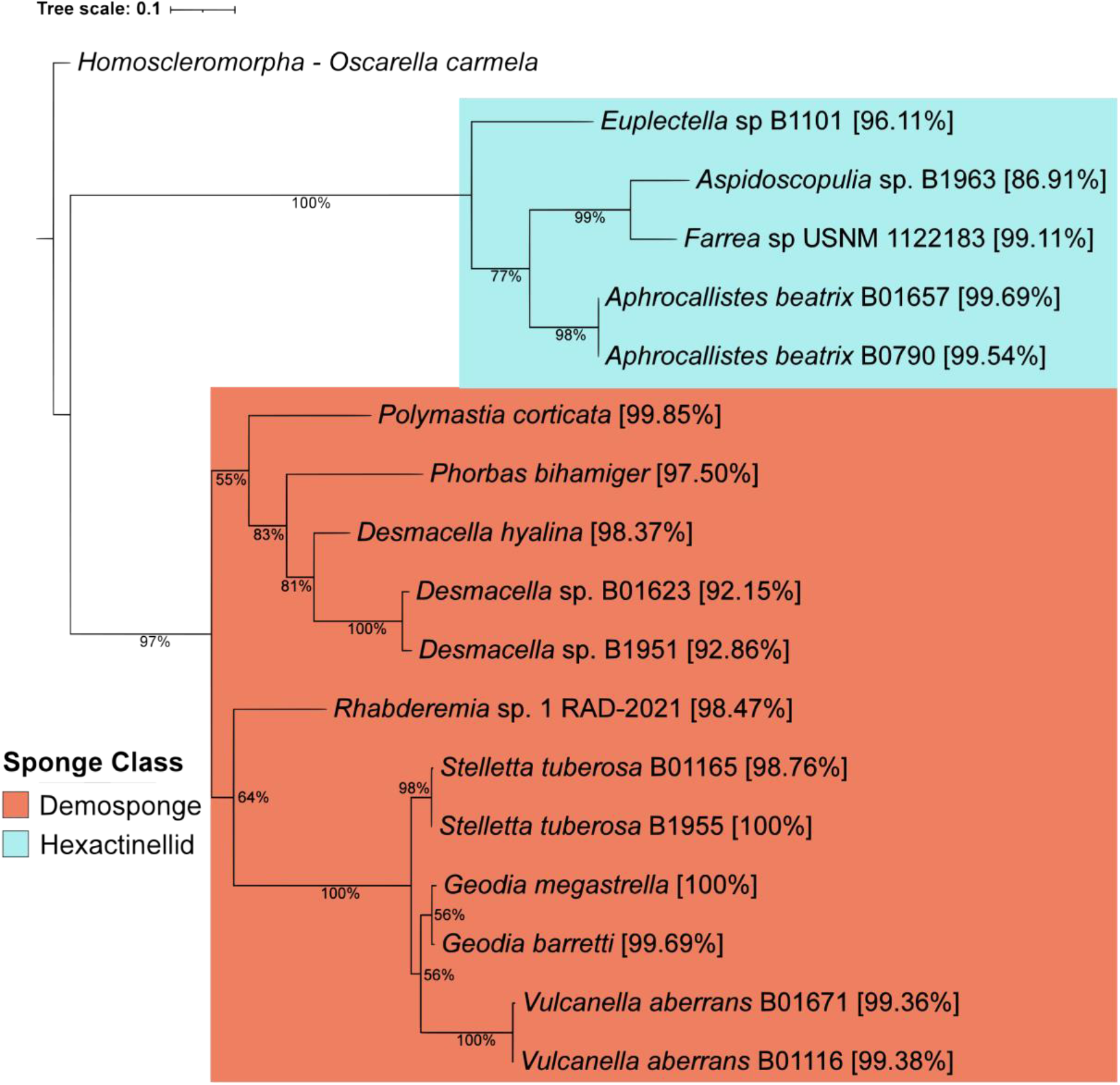
Maximum likelihood tree for 17 deep-sea sponge samples based on the mitochondrial cytochrome oxidase (COI) gene barcode. Bootstrap values are shown as a percentage of 1000 replications. Taxon branches are labelled with the closest relative from BLASTn (% ID) and the sampling site. Demospongia are highlighted in orange and Hexactinellida in blue. The slime sponge *Oscarella carmela* COI sequence is used as an outgroup.

### Microbial amplicon sequence variants (ASVs) and richness

Sequence abundance from the sponge microbiota for each sponge sample ranged from 200,223 to 437,147 with an average of 292,684 features per sponge. While individual sponges varied, across the whole dataset there was higher amount of archaeal (3,514,471) than bacterial abundance (3,185,734) and for *Hertwigia sp.* MD-2008 and Hexactinellid B01175 archaea represented over 90% of the reads (Supplementary Figure 1). Sequencing depth was sufficiently deep to be confident in recovering an accurate estimate of diversity, with all samples reaching at least 200,000 reads after DADA2 denoising (Supplementary Table 4) and approaching saturation in rarefaction curves (Figure 3a, Supplementary Figure 2).

**Figure 3.**
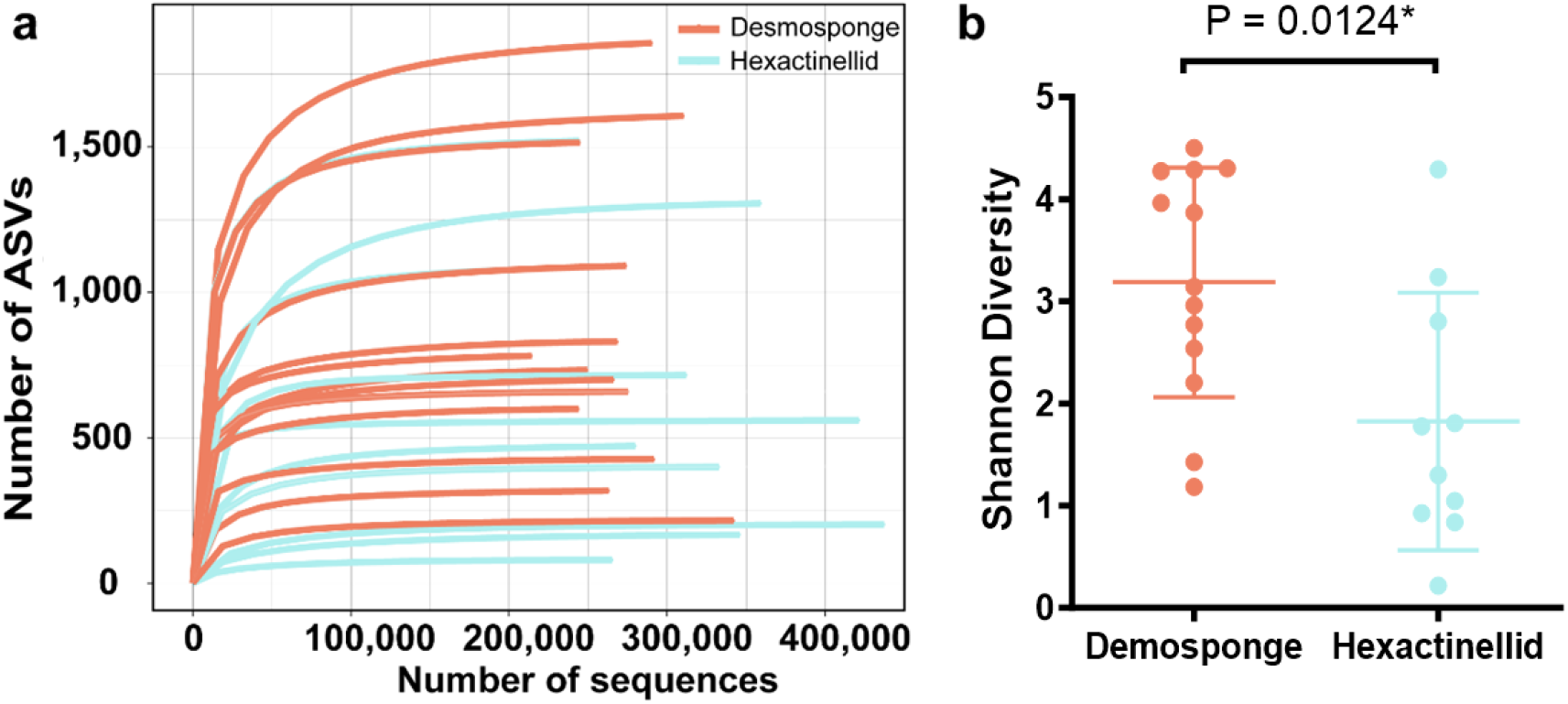
Microbial amplicon sequence variants (ASVs) and Shannon diversity derived from 23 deep-sea sponges. (**a**) Rarefaction curves of 16S rRNA gene diversity for deep-sea Atlantic sponge samples displayed by sponge class. Results from the hexactinellid *Euplectella* sp B1101 (1534 ASVs) and the demosponge *Vulcanella aberrans* B0116 (1530 ASVs) superimpose and are difficult to distinguish on this plot. (**b**) Shannon diversity associated with each sponge, presented by sponge class.

The total number of ASVs (*i.e.* richness) present across the 23 sponges was 11,465. The microbiota of sponge *Desmacella sp.* B1951 had the greatest number of ASVs with 1864, and this was over 20 times the number of ASVs in the least diverse sponge (*Aphrocallistes beatrix* B01657, 80 ASVs; Figure 3a, Supplementary Table 5). There was considerably higher bacterial richness at the Kingdom level with 10,844 ASVs assigned to Bacteria, 595 ASVs assigned to Archaea, and 26 ASVs that could not be assigned. Hexactinellid samples (*n* = 10) had 4698 bacterial and 315 archaeal ASVs, while Demosponge samples (*n* = 13) collectively had 6753 bacterial and 378 archaeal ASVs. Number of observed ASVs was slightly higher in demosponges compared to hexactinellids, but this difference was not statistically significant (p value = 0.298). Demosponges did, however, exhibit a significantly higher Shannon diversity index (p value = 0.0124) (Figure 3b). The range of ASVs within the dataset was large however, and certain hexactinellid samples from the *Euplectellidae* family (*Euplectella* sp. B1101 and *Hertwigia* sp. MD-2008) had amongst the highest number of ASVs (Supplementary Table 5). Examining the impact of depth on richness and diversity, no correlation was observed between increasing depth on observed ASVs (p value = 0.189414) or Shannon diversity (p value = 0.7624). However, a relationship emerged between depth and archaeal ASVs in demosponges, with a decrease in the number of observed archaea as depth increased, this pattern was exclusive to demosponges (p value = 0.002209).

### Taxonomic distribution and relative abundance of the sponge microbiota

We identified a total of 41 bacterial phyla and 4 archaeal phyla across the 23 sponge species analysed (Figure 4). There were 8 phyla found across all sponges in the dataset: Thermoproteota, Pseudomonadota, Chloroflexota, Desulfobacterota, Bacteroidota, Actinobacteriota, Gemmatimonadota and Myxococcota. Bdellovibrionota was present in low abundance in all samples except *Desmacella hyalina*. Acidobacteria, a dominant phylum in certain sponges, was present in all sponges except for *Aphrocallistes beatrix* B01657. There was a significant amount of reads unclassified even at the phylum level, this was particularly prominent in the two novel *Desmacella* sponge species, which had 56.11% (sp. B01623) and 32.16% (sp. B1951) of the relative abundance from unclassified reads. The diversity of phyla hosted varied across species, with *Aphrocallistes beatrix* B01657 hosting as few as 13 phyla, while *Vulcanella aberrans* B01671 exhibited a rich diversity of 38 phyla.

**Figure 4.**
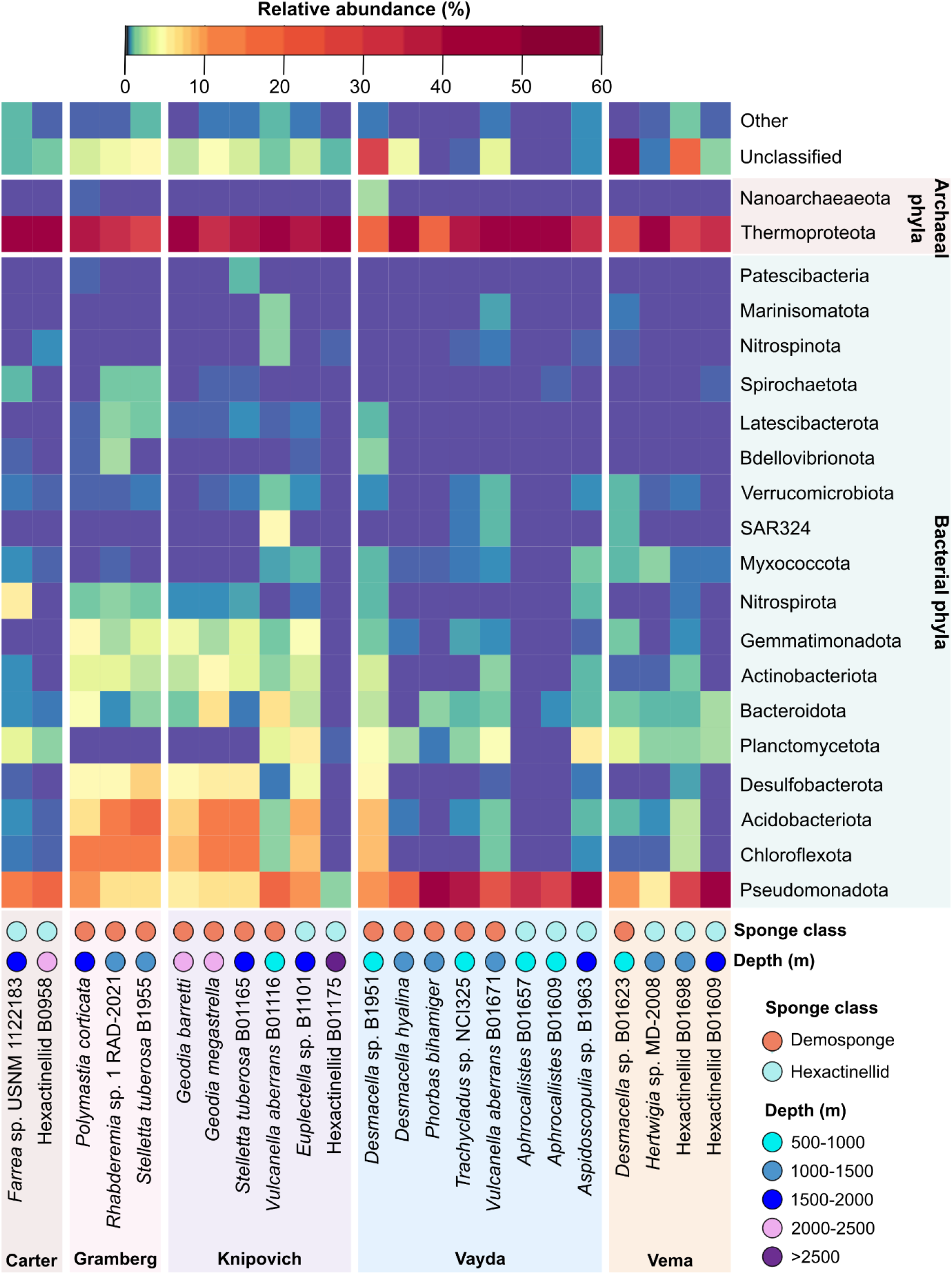
Abundant phyla in the deep-sea sponge microbiome, grouped by sampling location. Only phyla present with a relative abundance > 0.5% of total taxa in at least one sample are reported.

Archaea were more abundant than bacteria, with most archaeal abundance represented by phylum Thermoproteota (99.78%). This came almost completely from the *Nitrosopumilaceae* family (99.73%) which were by far the most abundant single taxa in our samples, where the second and third most abundant were family *CG1-02-33-12* (Nanoarchaeota phylum) and *Thalassarchaeaceae* (Thermoplasmatota phylum) with 0.11% and 0.04%. The most abundant bacteria were Pseudomonadota, particularly the classes Alpha-and Gammaproteobacteria, although Alphaproteobacteria organisms were lacking from both samples of *Aphrocallistes beatrix* obtained from the Vayda sea mount. The sponge associated Poribacteria phylum, thought to be primarily associated with sponges, were notably scarce among our deep-sea samples, but was identified in 5 demosponge samples and 2 hexactinellids. When present, this phylum comprised a tiny fraction of the overall diversity, falling beneath the >0.5% relative abundance threshold for plotting on Figure 4.

### Sponge species and geographic influence on the deep-sponge microbiome

To determine the influence of biotic and abiotic factors on the sponge microbiome, we performed Hellinger transformation on the ASV dataset and conducted a PERMANOVA analysis against three factors: sponge taxonomic class, sampling site, and sampling depth category. The results suggest that sponge class only explained 9% of variation in the microbiome, whereas sampling site and the depth category were responsible for 32% and 16% of the total variation, respectively (Table 1). Mantel tests showed significant correlation (*p*-value = 0.013) between the ASV dataset and sponge depth (*r* = 0.223) but not with geographic distances (*r* = 0.031, *p*-value = 0.302). These results show that while sampling site and depth impact microbiome and illustrate the important influence of locality on community structure (52).

**Table 1.**
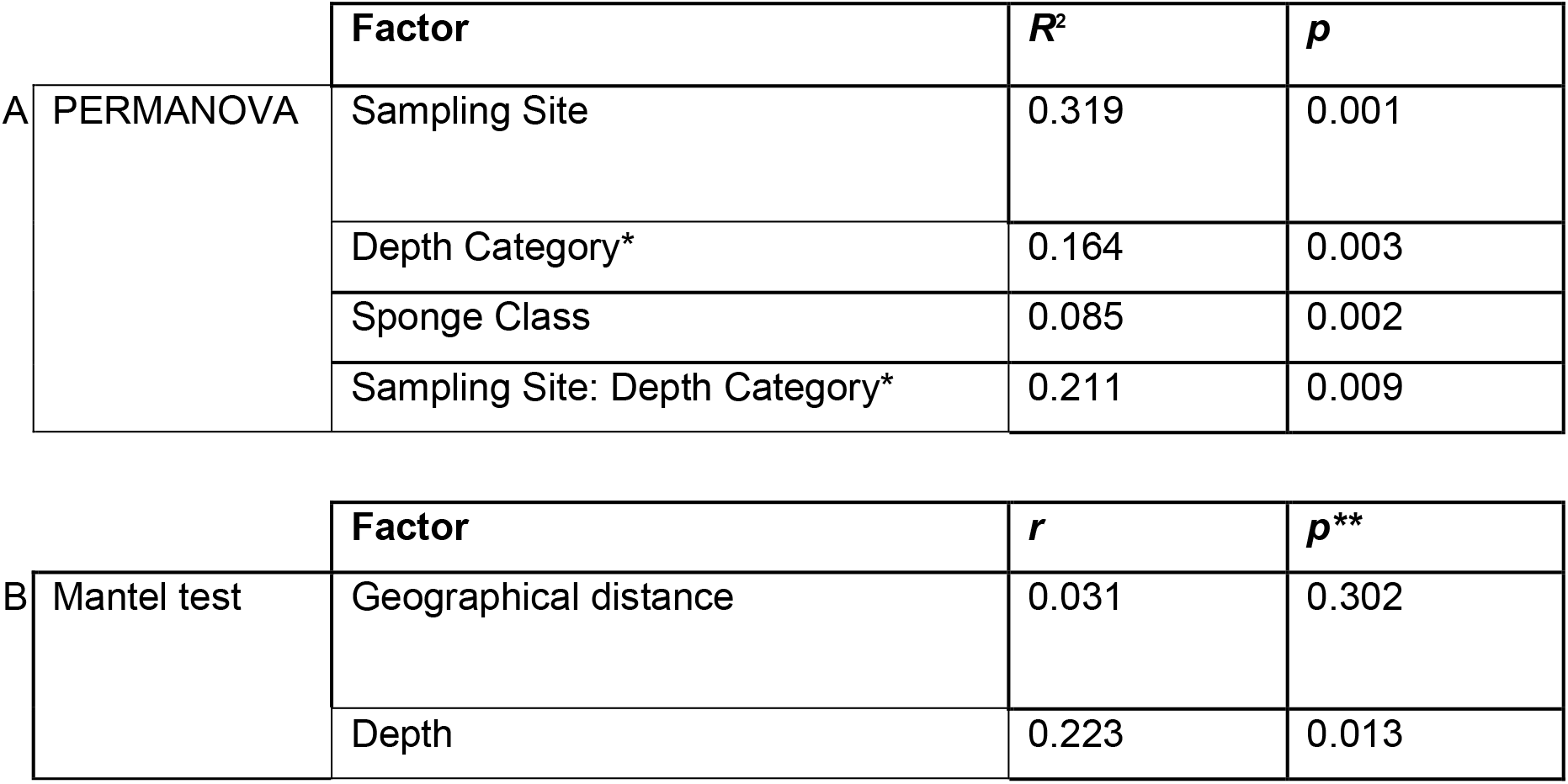
PERMANOVA (A) and Mantel (B) tests performed on Hellinger-transformed ASV dataset, using Bray-Curtis dissimilarity matrix with 1000 permutations. *Depth categories as in Figure 4. **Adjusted *p*-values using the Holm-Bonferroni method.

Next, we conducted a Non-Metric Multidimensional Scaling (NMDS) analysis of the community composition against the same factors of sponge class, site and depth (our dataset was insufficient for within-species comparisons). The NMDS groupings by sponge class were overlapping to some degree, although could be distinguished (Figure S5a). Sampling site and depth did not seem to be associated with clearly distinct clusters (Figure S5b-c). The most obvious individual cluster for sampling site was formed by the three demosponge samples from Gramberg Seamount (Figure S5b). Collectively, while PERMANOVA reveals that site and depth are generally important factors in explaining variation in microbial communities, our limited dataset does not show clear and obvious distinctions in community composition when comparing at the finer-grained level of sites or specific depths.

Finally, previous studies have suggested that sponge-associated microbial communities exhibit a high degree of host specificity at lower taxonomic ranks (11). We sought to investigate this phenomenon at the species level. To accomplish this, we performed a co-phylogenetic analysis relating the phylogenetic structure of sponge microbial communities compared to the COI gene-based phylogenetic tree from the host sponges. This comparative analysis showed similar, but not identical clading between the microbial assemblage and the host phylogeny (Figure 5). Our findings highlight the predominance of host species in shaping the structure of the deep-sea sponge microbiome, mostly irrespective of variations in depth or geographical distribution of the same sponge species. One exception here is the two samples of *Stelletta tuberosa*, where it’s clear variation in depth and site play a larger role. Additionally, an anomaly was noted in the sponge *Euplectella* sp. B1101, which demonstrated a phylogenetic affinity more closely aligned with *Geodia* sponges than with other members of the Hexactinellida.

**Figure 5.**
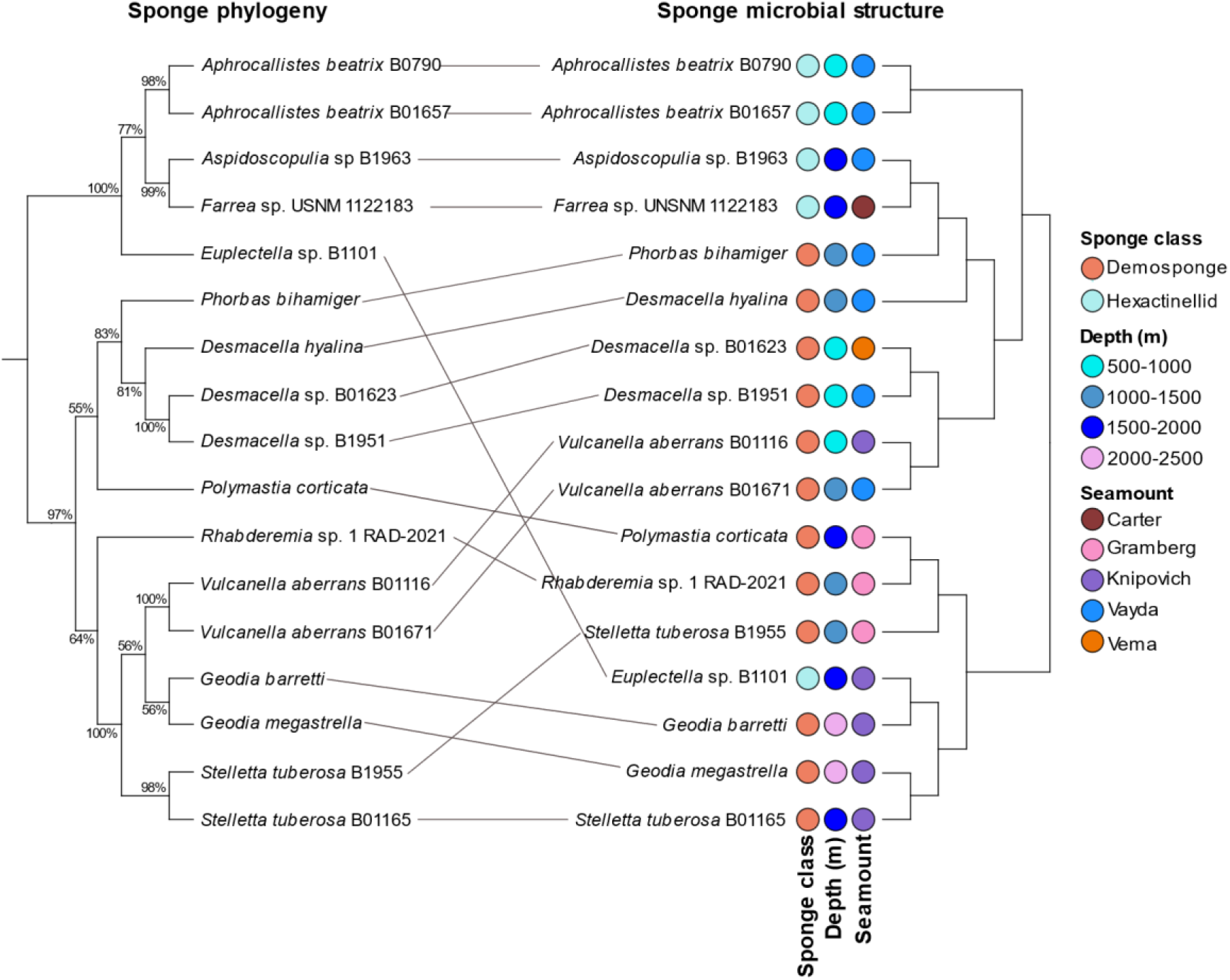
The association between sponge host phylogeny based on COI gene sequences and associated microbial community structure. The left tree represents the COI gene-based phylogenetic relationships among sampled sponge species, while the right tree illustrates the phylogenetic structure of their respective microbial communities. Lines connecting the two trees highlight the observed congruence, demonstrating the influence of host phylogeny on microbial assemblage.

## Discussion

The microbial ecology of the deep-sea remains underexplored. This study employs metataxonomic methods to investigate the microbial community structure of 23 sponges from 5 Atlantic seamounts, including many sponges that have not previously been characterised and are likely to be unique to the deep ocean. We reveal a microbiome predominantly consisting of ammonia-oxidizing archaea from the family *Nitrosopumilaceae*. Importantly, our data provides further evidence that host-sponge phylogeny is associated with microbiome composition, contributing to a further understanding of deep-sea sponge microbial ecology.

### Thermoproteota dominate the sponge microbiome

Our findings further contribute to the understanding of deep-sea sponge microbiota, corroborating prior observations of high abundance of archaea and particularly dominance of Thermoproteota (6, 12-14). Metagenome-assembled genomes of sponge associated *Nitrosopumilaceae* show a high degree of symbiotic adaption (53). In our study an overwhelming 99.75% of archaeal reads were identified as Thermoproteota, underscoring their key role to oxidise ammonia, and so contribute to nitrogen fixation (54, 55).

### Methodological Limitations

The absence of seawater controls limits the extent to which this study can be compared to others. Though we implemented stringent rinsing protocols, seawater contamination remains a concern as do reads present in the blank extraction. The sponges studied here are a subset of a larger collection collected to study biomineralization (56) and subsequently bioprospecting (24, 57, 58). The samples thus represent an unbiased general survey, with a modest sample size, rather than a systematic effort to understand a particular sponge species or geographical location, for this analysis we direct readers to large systematic studies of the deep-sea sponge microbiome (11, 12).

### Comparison with large scale studies

Our findings align with Busch et al.’s extensive study of 1077 deep-sea sponges (11), but add novel, previously unstudied genera of deep-sea sponges. The use of GreenGenes2 reference tree improves microbial taxonomic assignment and facilitating comparability with the Genome Taxonomy Database (19, 59). Our study also confirms the presence of both LMA and HMA sponges in deep-sea environments (60). Interestingly, while hexactinellids generally appear to be LMA, we do identify a potential HMA Hexactinellid class sponge from the genus *Euplectella.* a genus that has not been characterised previously and has a microbiome similar to phylogenetically distance sponges. Busch et al. noted that *Amphidiscella caledonica* (family *Euplectellidae*) diverged from typical LMA glass sponges, showing high Chloroflexota abundance and several other members of *Euplectellidae* family clustered with more with demosponges. We identify two further *Euplectellidae* genera (*Euplectella, Hertwigia*) which deviate from the classical composition of LMA glass sponges.

### Genus-Specific Comparisons

Our study also includes two representatives of the demosponge genus *Geodia* collected from the Knipovich seamount. Both sponges have similar community profiles to samples of *Geodia hentscheli* and *Geodia barretti* from other studies (6, 16, 61), being dominated by Pseudomonadota, Chloroflexota and Acidobacteriota, as well as Desulfobacterota and Gemmatimonadota. Although these other studies did report a higher abundance of Poribacteria that was not observed here. Otherwise, samples of *Stelletta tuberosa* from Gramberg and Knipovich seamounts had relatively dissimilar profiles based on the tanglegram analysis, whereas the two *Vulcanella aberrans* samples from Knipovich and Vayda sites clustered together strongly. The relative dominance of Chloroflexota and Acidobacteriota in *Stelletta tuberosa* was previously observed in another member of the genus, *Stelletta normani* obtained from canyons of the North Atlantic and is consistent with the assignment of this species as a deep-sea HMA sponge (13).

### Influence of sponge host and biogeographical factors

Deep-sea sponge microbial communities are influenced by the sponge host species and biogeographical factors such as nutrient profile and the surrounding seawater (16). Our study emphasises the role of depth in shaping sponge associated communities, a finding which is supported by Busch et al. (11), the *Tara* Oceans project (62) but contrasting with other systemic studies (12). A study of seamounts impact on the microbial composition of sponges did highlight the impact of depth on phyla distribution (16), suggesting depth could impact communities in an environment specific manner. The deep-sea sponge microbiome project highlighted a weak but significant biogeographical impact on community composition, albeit over a much larger distance of 10,000 km (11). In our study, significant differences between sponges at various sites suggest that isolation by seamounts, rather than geographical distance, influences community structure. Influence of sampling site over geographical distance has been previously observed in deep-sea microbiomes both in sponge hosts and deep-sea sediments (52, 63).

We also identified accordance between sponge host phylogeny and sponge microbiome structure (Figure 5), indicating host-driven microbiome composition. For example, the microbiomes of two *Aphrocallistes beatrix* sponges, both collected from the Vayda seamount, displayed a consistent microbial community, also supporting previous findings from the same species (12). The identification of a co-phylogenetic relationship, known as phylosymbiosis, has been previously identified in coral (64), animal (65) and shallow reef sponge microbiomes (66). While our findings indicate an influence of host phylogeny on microbiome structure, this relationship is not absolute, reflecting that deep-sea sponges may prioritize functional redundancy over species-specificity in their microbial associations. This is supported by a comparative study of 39 HMA sponge species, revealing a more pronounced species-specificity in microbial composition among shallow water sponges than in their deep-sea counterparts (67).

## Conclusion

Overall, our study explores the microbiomes of previously unstudied deep-sea sponges and identify depth and sponge phylogeny to be drivers of microbial composition. Our results particularly highlight distinct microbiomes in the *Euplectellidae* family, differing from other glass sponges which warrant further investigation.

## Author Contributions

S.E.W. set out the methodology of the study, performed data collection, conducted bioinformatic data curation and initial investigations of the data. S.E.W. and G.V. conducted data analysis and produced visualizations for the manuscript. S.E.W. wrote the original draft of the manuscript with help from G.V., P.C., M.L., J.E.M and P.R.R. All authors reviewed and edited the manuscript. P.C. and P.R.R. conceived the project, provided supervision and acquired funding. All authors have read and agreed to the published version of the manuscript.

## Acknowledgments

The authors thank the Bristol Genomics Facility for performing the Illumina sequencing. We thank Laura Robinson, Chief Scientist for expedition JC094, and Kate Hendry for initial collection of the sponge samples and facilitating subsequent access.

## Supplementary materials

**Figure S1.**
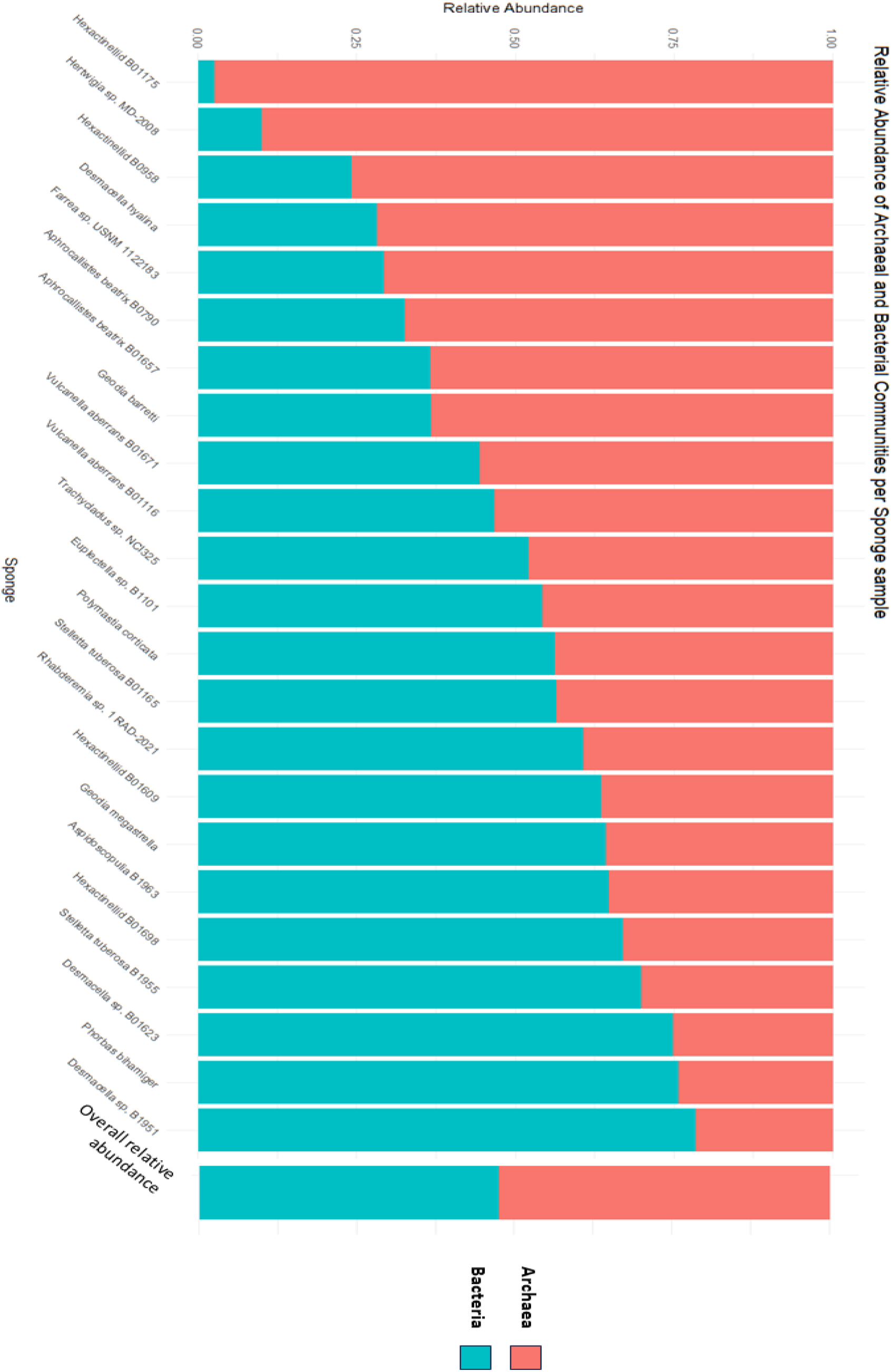
Relative abundance of Archaea and Bacteria in the deep-sea sponge microbiome. Overall relative abundance is taken from the entire dataset.

**Figure S2.**
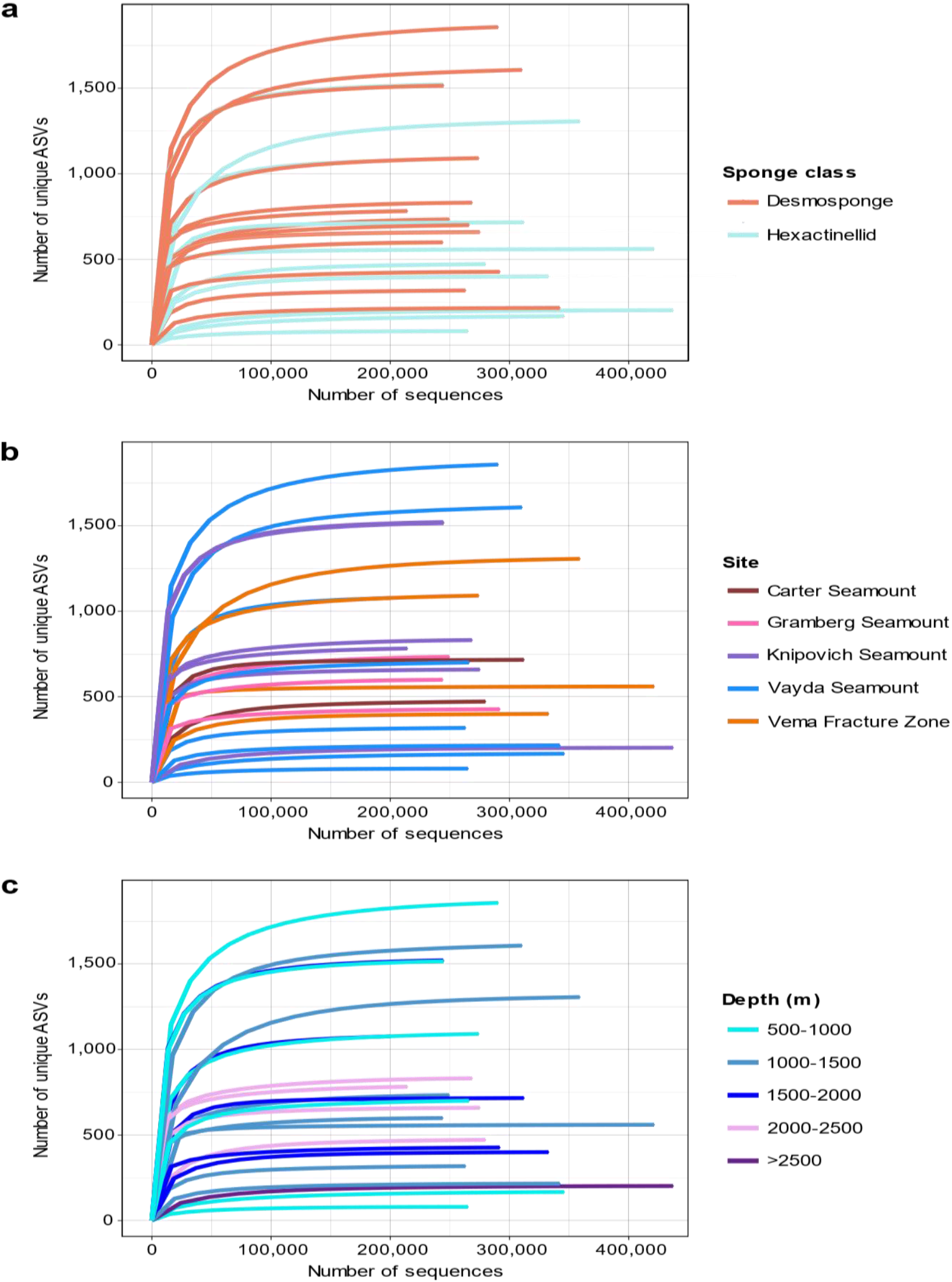
Rarefaction curves of 16S rRNA gene diversity for deep-sea Atlantic sponge samples. Displayed by a) sponge class b) site and c) depth.

**Figure S3.**
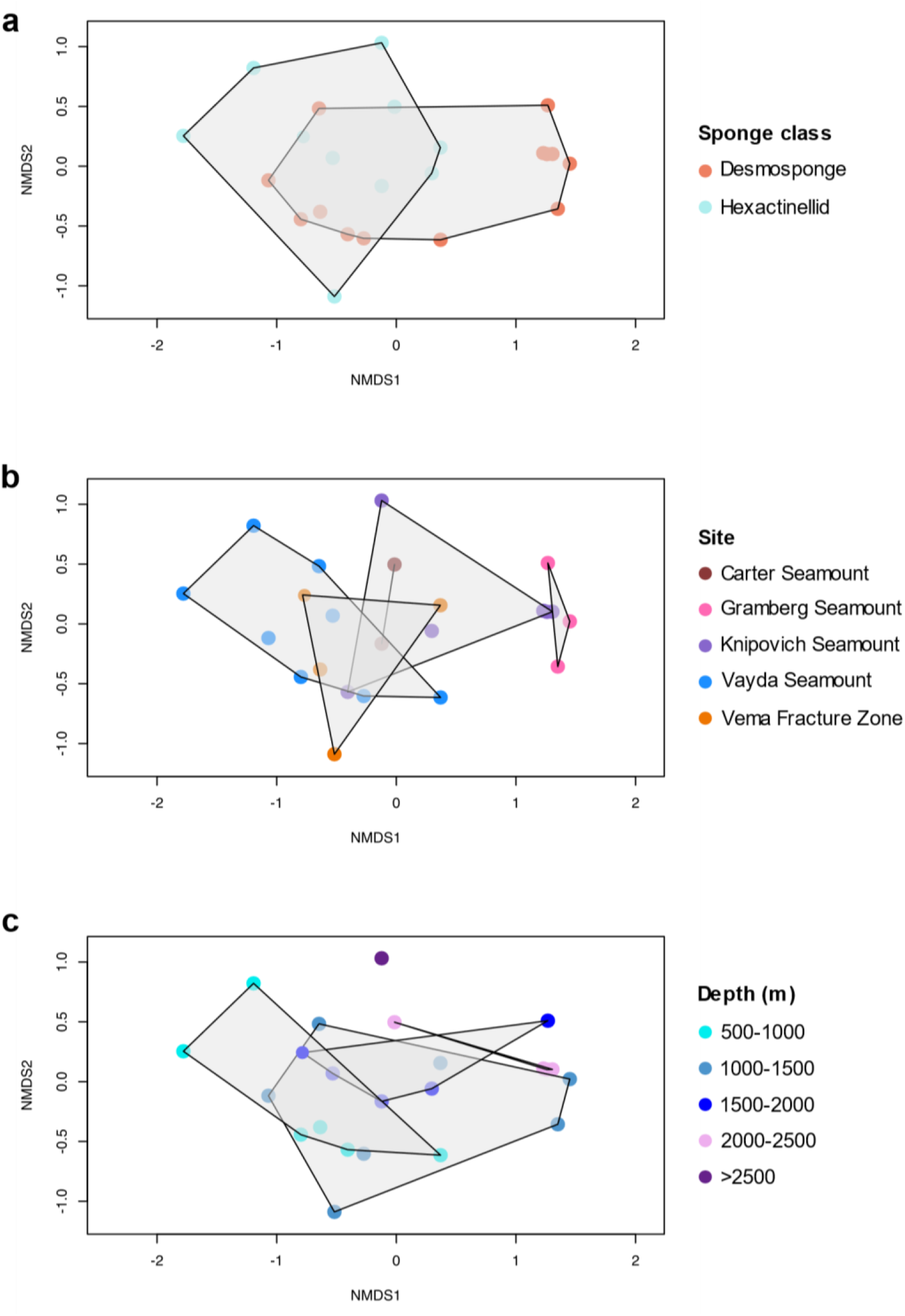
Compositional similarity of the deep-sea sponge microbiome in the Atlantic Ocean. NMDS performed on the Hellinger-transformed ASV dataset. Samples are grouped by (a) sponge class, (b) sampling site and (c) sampling depth.

**Table S1.**
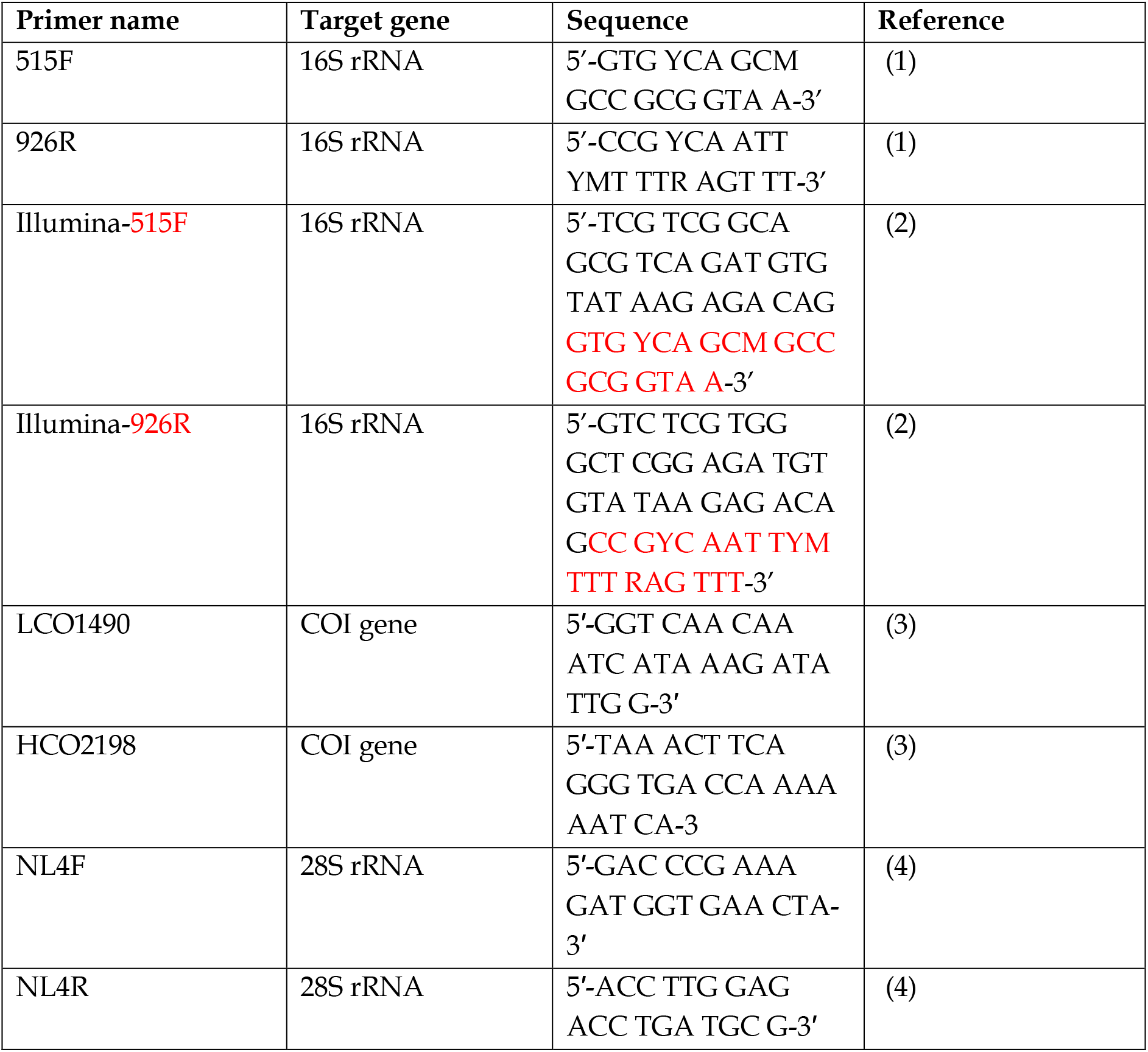
PCR primer sequences used in the study including the target gene and the nucleotide sequence.

**Table S2.**
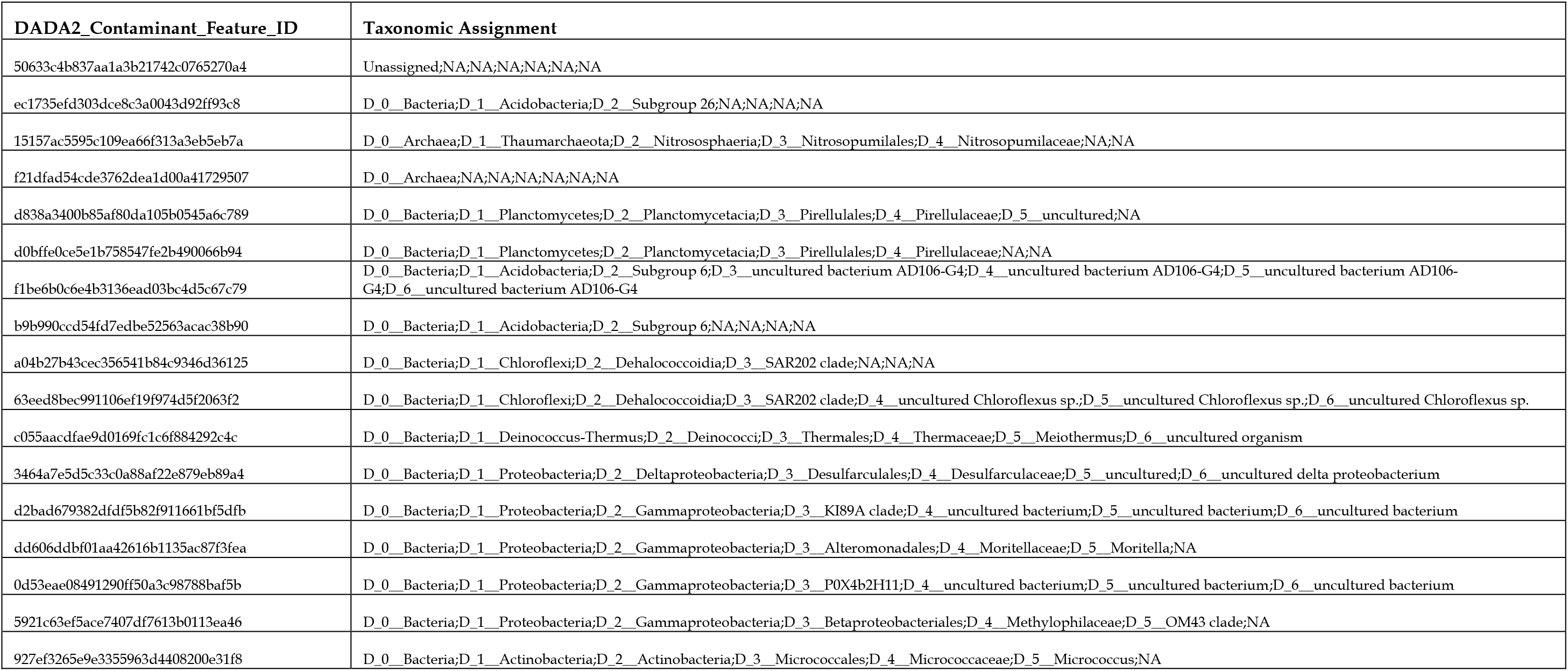
DADA2 ASV features identified by decontam 1.8.0 as contaminants which were then removed from the final data set.

**Table S3.**
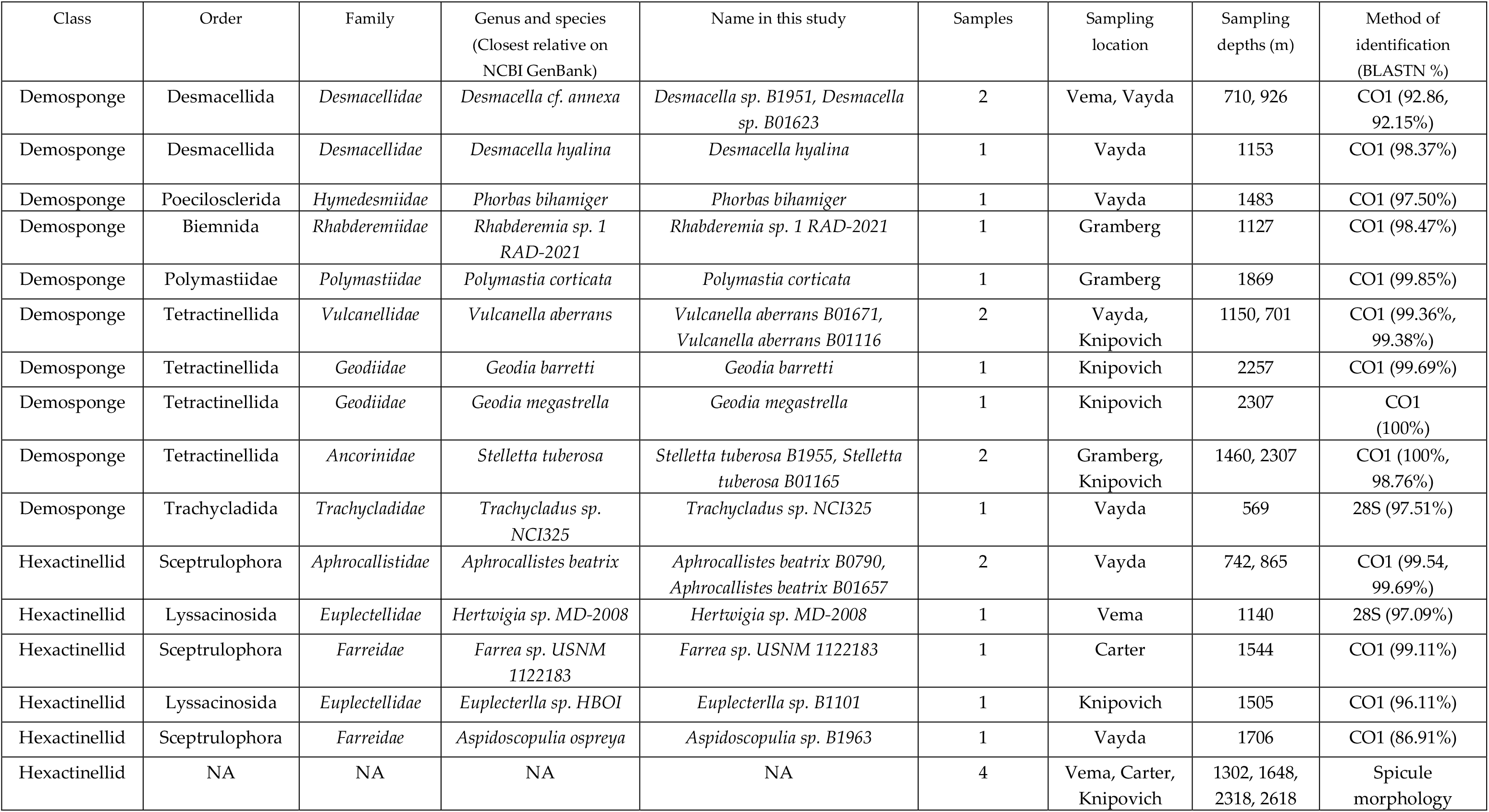
Taxonomic assignment of sponges, sampling site and depth information. Method of identification and BLASTn % identity to closest relative.

**Table S4.** DADA2 denoising of raw sequencing to amplicon sequence variants for each sponge sample and two negative controls.

| *Sample-code* | *Sample-ID* | *input* | *filtered* | *denoised* | *merged* | *non-chimeric* |
| --- | --- | --- | --- | --- | --- | --- |
| B0958 | Hexactinellid B0958 | 362870 | 306353 | 306353 | 293968 | 279823 |
| B0507 | *Hertwigia* sp. MD-2008 | 454548 | 402398 | 402398 | 392163 | 359254 |
| B01671 | *Vulcanella aberrans* B01671 | 451350 | 370365 | 370365 | 341796 | 310478 |
| B01165 | *Stelletta tuberosa* B01165 | 411178 | 355792 | 355792 | 303745 | 214299 |
| B1101 | *Euplectella* sp. B1101 | 346965 | 281666 | 281666 | 262303 | 244673 |
| B01637 | *Desmacella hyalina* | 326105 | 276846 | 276846 | 271328 | 263138 |
| B0768 | *Farrea* sp. USNM 1122183 | 403221 | 326306 | 326306 | 323577 | 312162 |
| B01623 | *Desmacella* sp. B01623 | 408675 | 353907 | 353907 | 328635 | 274158 |
| B01171 | *Geodia barretti* | 397309 | 334193 | 334193 | 300019 | 275125 |
| B1951 | *Desmacella* sp. B1951 | 446100 | 370664 | 370664 | 327932 | 291343 |
| B1963 | *Aspidoscopulia* sp. B1963 | 317991 | 256963 | 256963 | 248992 | 200223 |
| B01657 | *Aphrocallistes beatrix* B01657 | 304879 | 272175 | 272175 | 269622 | 265216 |
| B01116 | *Vulcanella aberrans* B01116 | 433479 | 356167 | 356167 | 331611 | 244735 |
| B0790 | *Aphrocallistes beatrix* B0790 | 419338 | 362530 | 362530 | 354738 | 345848 |
| B01683 | *Phorbas bihamiger* | 455059 | 358571 | 358571 | 355041 | 342180 |
| B01609 | Hexactinellid B01609 | 424806 | 375249 | 375249 | 370658 | 332808 |
| B01175 | Hexactinellid B01175 | 512952 | 456317 | 456317 | 454292 | 437157 |
| B01686 | *Trachycladus* sp. NCI325 | 402187 | 334188 | 334188 | 317666 | 266038 |
| B1954 | *Rhabderemia* sp. 1 RAD-2021 | 400168 | 345122 | 345122 | 288800 | 251398 |
| B1955 | *Stelletta tuberosa* B1955 | 474722 | 374922 | 374922 | 309185 | 244069 |
| B01140 | *Geodia megastrella* | 498304 | 415785 | 415785 | 349302 | 268611 |
| B01698 | Hexactinellid B01698 | 564951 | 435208 | 435208 | 432029 | 423233 |
| B1983 | *Polymastia corticata* | 401621 | 346304 | 346304 | 319266 | 291842 |
| Blank | - | 429850 | 364245 | 364245 | 355614 | 350173 |
| Gel-extract-neg | - | 2269 | 662 | 662 | 462 | 398 |

**Table S5.** Alpha diversity metrics of each deep-sea sponge sample ordered by Observed ASVs.

| Sponge sample | Observed | Chao1 | Shannon | InvSimpson |
| --- | --- | --- | --- | --- |
| *Desmacella* sp. B1951 | 1864 | 1887.90 | 4.50 | 11.35 |
| *Vulcanella aberrans* B01671 | 1610 | 1644.71 | 2.77 | 3.55 |
| *Euplectella* sp. B1101 | 1534 | 1552.55 | 4.29 | 7.97 |
| *Vulcanella aberrans* B01116 | 1530 | 1537.48 | 4.29 | 9.52 |
| *Hertwigia* sp. MD 2008 | 1271 | 1294.95 | 1.30 | 1.44 |
| *Desmacella* sp. B01623 | 1098 | 1109.23 | 2.96 | 3.32 |
| *Aspidoscopulia* sp. B1963 | 1086 | 1102.60 | 2.80 | 4.33 |
| *Geodia megastrella* | 851 | 853.76 | 4.30 | 19.17 |
| *Stelletta tuberosa* B01165 | 784 | 788.36 | 4.28 | 16.75 |
| *Rhabderemia* sp. 1 RAD 2021 | 739 | 741.25 | 3.87 | 6.80 |
| *Farrea* sp USNM 1122183 | 734 | 736.15 | 1.81 | 2.13 |
| *Trachycladus* sp. NCI325 | 706 | 710.50 | 2.20 | 3.35 |
| *Geodia barretti* | 669 | 670.06 | 2.54 | 2.63 |
| *Stelletta tuberosa* B1955 | 595 | 595.61 | 3.96 | 10.67 |
| Hexactinellid B01698 | 566 | 568.14 | 3.24 | 8.48 |
| Hexactinellid B0958 | 481 | 495.44 | 1.05 | 1.69 |
| *Polymastia corticata* | 439 | 440.00 | 3.14 | 5.34 |
| Hexactinellid B01609 | 411 | 412.27 | 1.78 | 3.19 |
| *Desmacella hyalina* | 312 | 315.06 | 1.18 | 1.92 |
| *Phorbas bihamiger* | 216 | 218.00 | 1.43 | 3.22 |
| Hexactinellid B01175 | 202 | 202.79 | 0.22 | 1.06 |
| *Aphrocallistes beatrix* B0790 | 150 | 151.00 | 0.93 | 1.94 |
| *Aphrocallistes beatrix* B01657 | 80 | 80.43 | 0.84 | 1.99 |

